# Species-Specific Responses of Bird Song Output in the Presence of Drones

**DOI:** 10.1101/2020.07.19.211045

**Authors:** Andrew M. Wilson, Kenneth S. Boyle, Jennifer L. Gilmore, Cody J. Kiefer, Matthew F. Walker

## Abstract

Drones are now widely used to study wildlife, but applications for studying bioacoustics have been limited. Drones can be used to collect data on bird vocalizations, but an ongoing concern is that noise from the drones could change bird vocalization behavior. To test this behavioral impact we conducted an experiment using 30 sound localization arrays to track the song output of seven songbird species before, during, and after a 3-minute flight of a small quadcopter drone hovering at 50 m above ground level. We analyzed 8,303 song bouts, of which 2,285 song bouts of 184 individual birds were within 50 meters of the array centers. We used linear mixed effect models to assess patterns in song output showed patterns that could be attributed to the drone’s presence. We found no evidence of any effect of the drone for five species: American Robin *Turdus migratorius*, Common Yellowthroat *Geothlypis trichas*, Field Sparrow *Spizella pusilla*, Song Sparrow *Melospiza melodia*, and Indigo Bunting *Passerina cyanea*. However, we found a substantial decrease in Yellow Warbler *Setophaga petechia* song detections during the 3-minute drone hover, such that there was an 81% drop in detections in the 3^rd^ minute (Wald-test, p<0.001), compared with before the drone’s introduction. In contrast, the number of singing Northern Cardinal *Cardinalis cardinalis* increased after the drone was introduced, and remained almost five-fold higher for 4-minutes after the drone departed (P<0.001). Further, we found an increase in cardinal contact/alarm calls when the drone was overhead, with the elevated calling-rate sustaining for 2 minutes after the drone had departed (P<0.001). Our study suggests that responses of songbirds to drones may be species-specific, an important consideration when proposing the use of drones in avian studies. We note that recent advances in drone technology have resulted in much quieter drones, which makes us hopeful that the impacts that we detected could be greatly reduced.

## INTRODUCTION

Drones or Unoccupied Aircraft Systems (UASs) are now well-established tools in field ecology, especially for purposes of mapping habitats or counting and tracking megafauna (Vermeulen *et al*., 2013; Christiansen *et al*., 2016; Witczuk *et al*., 2017; Kelaher *et al*., 2020). Drones have been proven to be effective tools for conducting aerial surveys large birds, especially those found in open habitats (Ratcliffe *et al*., 2015; Barnas *et al*., 2019) or that nest or roost in tree canopies (Weissensteiner, Poelstra and Wolf, 2015; Afán, Máñez and Díaz-Delgado, 2018), but there have been few studies that show drones to be useful for monitoring small birds, or those that inhabit dense vegetation, where the reach of aerial imagery is limited. Despite some of the potential advantages of drones, for example their efficiency, ease of access, and potential to reduce disturbance (Borrelle and Fletcher, 2017), drones are not widely used for monitoring smaller birds, such as songbirds, which constitute the majority of birds species and abundance in most terrestrial environments (Musgrove *et al*., 2013; Inger *et al*., 2015; Rosenberg *et al*., 2019). Recent advances coupling drones with audio-recorders for bioacoustics monitoring (Wilson, Barr and Zagorski, 2017; Kloepper and Kinniry, 2018), and infrared cameras to detect passerine nests (Scholten *et al*., 2019), suggest that the use of drones for monitoring of smaller birds could grow.

An important drawback of drones is that they emit noise, which, in addition to their visual impacts, can disturb wildlife. There have now been many studies that investigate the impacts of drones on wildlife. A review of these concluded that smaller drones, flown at greater heights, and spending reduced time overhead, were less likely to elicit a response in study organisms (Mulero-Pázmány *et al*., 2017). However, there are still many unknown impacts of drones that have yet to be studied, especially for non-target organisms and their responses.

In developing protocols to use drones for bioacoustics monitoring, we are interested in the possible impacts of drones on songbirds for two important reasons: firstly, if drones do impact songbird behavior, it introduces an ethical decision into whether they should be used at all, and secondly, if drones cause songbirds to move locations or change vocalization behavior, data derived from airborne bioacoustics could be biased. In this study, we aim to assess the impacts of hovering a drone over a study area for the intention of airborne bioacoustics monitoring, to test the hypothesis that drones initiate vocal behavior responses in songbirds. While the bounds of our study are narrow--we focus on the impacts of a single drone model, flown at a specific height--we hope that this study sheds light on potential impacts of drones on songbirds, which to most ecologists using drones would be considered non-target species. Further, there is a wider concern that as an additional source of anthropogenic noise pollution, drones flown by recreationists as well as professionals may be adding to growing noise pollution problems. Our study will inform this broader context by assessing the ecological impact of noise pollution from drones on bird singing behavior.

## METHODS

To test to see whether there was a change in songbird vocalizing behavior that could be attributed to the drone’s presence, we set up an experiment whereby ambient bird sound in the area under the drone was recorded before, during and after a drone flight. Traditionally, bird population surveys are completed using a point count technique, in which ornithologists record visual and auditory bird cues for a fixed period of time at a series of pre-determined locations. Following a previous study (Wilson, Barr and Zagorski, 2017), we simulated an airborne bird point count by hovering a small quadcopter (DJI Mavic Pro) over a pre-determined count location for 3-minutes. Traditional point count duration is typically from 3 to 10 minute; we chose the lower end of that range because this minimizes flight time, thereby allowing for greater survey efficiency—potentially allowing for multiple points to be surveyed on each drone battery. Additionally, shorter flight duration is less likely to cause excessive disturbance (Mulero-Pázmány *et al*., 2017).

### Field Study

We used sound localization to locate singing birds in the vicinity of a ground-based Automated Recording Unit (ARU) array to track bird song output before, during, and after our drone flights (Figure 1i). By arranging four ARUs in a 50 × 50 m array, we were able to measure the differences in the time taken for each bird vocalization to reach each recorder, a method called “time difference of arrival” (TDOA). The differences in the time of arrival allowed us to triangulate birds in two dimensional space inside and beyond the array using the Sound Finder Microsoft Excel application (Wilson *et al*., 2014). The Sound Finder method has been proven to allow accurate localizations, with a mean error of 4.3 m, and 74% to within 10 m (Wilson *et al*., 2014).

**Figure 1.**
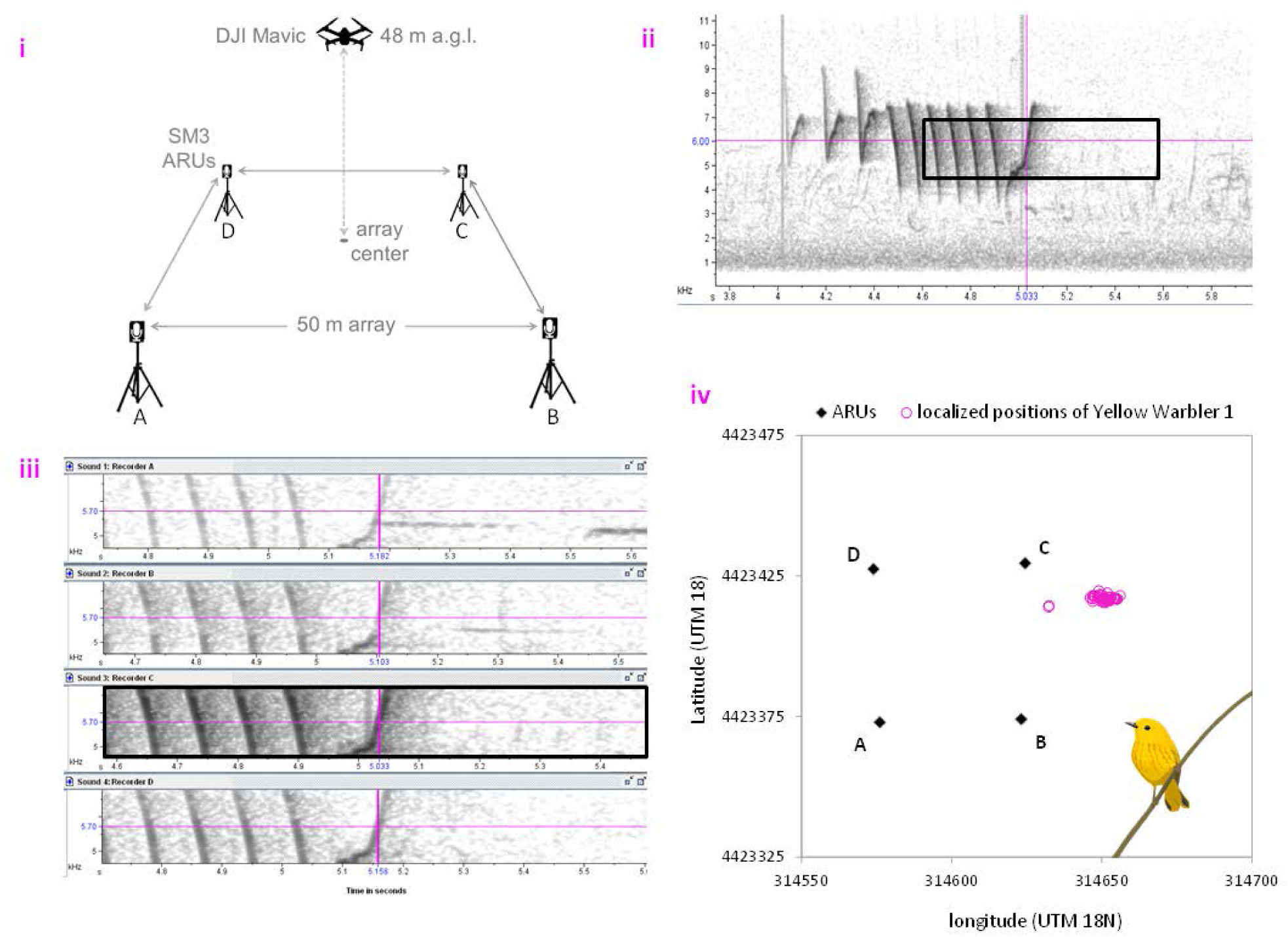
Methods. Pane **i** shows the setup of each experimental array, **ii** shows the spectrogram of a Yellow Warbler song from recorder C with a black rectangle showing inset in pane **iii**, which shows the measurement of time difference of arrival in Program Raven, and **iv** the estimated positions of the Yellow Warbler over the entire 11-minute experiment, relative to ARUs A, B, C and D.

We conducted the experiments in June and early July of 2017 at Pennsylvania State Game Lands 249, Heidlersburg, Adams County, Pennsylvania (39°56’14.6”N 77°10’38.6”W). The experiment had 30 replicates at locations on a 200 m grid to ensure a representative sample of the habitat and to reduce the likelihood of resampling birds between replicates. At each location, we deployed four SM3 ARUs (Wildlife Acoustics) on tripods 1.5 m off the ground in a 50 m x 50 m array centered on the predetermined grid points. The recorders were fitted with Garmin GPS time-synch receivers to synchronize their clocks. Once the recorders were set up, we retreated to at least 150 m from the center of the array and waited at least 10 minutes to allow normal bird activity to resume following possible human disturbance. After the resettlement period we deployed the drone from a base station at least 150 m away and programmed it to autonomously hover 48 m above ground level at the center of the array for three minutes, to simulate an aerial point count as described in Wilson et al. (2017). After returning the drone to the base station, we continued recording ambient noise in the study array for an additional five minutes. All flights were preprogrammed using the Litchi App (VC Technology Ltd.) to ensure high degrees of spatial precision.

### Bioacoustic Analysis and Sound Localization

We used Audacity to clip the recordings from each of the four ARUs to the exact same 11-minute time-span (to nearest millisecond), which included 4-minutes before the drone flight, 3-minutes while the drone was overhead, and 4-minutes after the drone flight. We used 4-minutes before and after the drone flight so that we could exclude the one-minute immediately before and after the drone-flight, during which the drone was audible on the recordings as it approached and departed the array. This allowed us to isolate 3 × 3-minute blocks for “before” (minutes 1-3) “during” (minutes 5-7) and “after” (minutes 9-11) the drone flight.

We focused our data analysis on seven species that were known to be present on the site in moderate to high densities: American Robin *Turdus migratorius*, Yellow Warbler *Setophaga petechia*, Common Yellowthroat *Geothlypis trichas*, Field Sparrow *Spizella pusilla*, Song Sparrow *Melospiza melodia*, Indigo Bunting *Passerina cyanea*, and Northern Cardinal *Cardinalis cardinalis*. Some other abundant species were not included in the study because we could not reliably identify individual birds in the audio recordings due to incessant song (Gray Catbird *Dumetella carolinensis*) and localized high densities (Red-winged Blackbird *Agelaius phoeniceus*). For our seven study species, we identified all song bouts that were clearly visible in spectrograms from all four ARUs (Figure 1ii). In a given recording, we often identified more than one individual of a given species and more than one song bout of a given individual. To connect the song bouts from the same individual, we assigned a unique identification code to identify each individual bird based on location, song volume (visually assessed in spectrogram), song-bout spacing, and unique characteristics of song.

Time differences of arrival were measured manually from spectrograms in program Raven Pro 1.5 (Figure 1iii). Spectrogram settings were Hanning window with 512 samples and 89% overlap. The time differences and air temperature at the time of the experiment (logged by the ARUs) were then input into the Sound Finder spreadsheet to localize each bird, with locations given in UTM coordinates. Each songbird location was added to a scatterplot in the Excel spreadsheet relative to the coordinates of the ARUs, to manually verify that the calculated coordinates were logical (Figure 1iv), for example, if the vocalization reached ARU C first, then the bird should be closer to that recorder than the others. Each individual bird was localized for every song bout throughout the 11-minute experiment, so that the bird’s movement and song output could be tracked through space and time.

While listening to the recordings to locate song bouts of our target species, we observed an apparent increase in Northern Cardinal “tik” contact/alarm calls when the drone was overhead. Because these calls are quieter than songs, and therefore difficult to localize because many are not detected on all four ARUs, we used a different approach to determine whether the rates of these calls changed in the presence of the drone. We used Kaleidoscope software (Wildlife Acoustics, 2019) to automatically detect Northern Cardinal “tik” calls in recordings from a single ARU (A) from each array. To train Kaleidoscope we developed a training set of 10 recordings of “tik” calls. All identified calls were verified aurally, to exclude calls of other species.

### Statistical Analysis

To test whether birdsong detections varied by time, we constructed linear mixed-effect models (GLMMs) using the ‘glmer’ function in R package lme4 (Bates *et al*., 2015). We conducted two sets of analyses 1) the probability of detected an individual bird in each minute and 2) the singing rate in each minute. For the first analysis, the response variable was whether or not an individual bird was detected, modelled with a binomial error distribution. We constructed eight models: a null model with individual bird as a random effect (model 1), a model with distance to bird from center of the array as a covariate (model 2), three models with time effects (models 3-5), and three models with time effects plus distance (models 6-8). The three time effects were: individual minutes (models 3 and 6); the 3-minute periods “before”, “during” and “after” (models 4 and 7); and “before” and “after” but with effects varying by minute during the drone flight (models 5 and 8; Figure 2). All eight models included individual bird as a random effect. Models were compared using AIC, the size of effects determined using odd-ratios, and significance determined using Wald tests.

**Figure 2.**
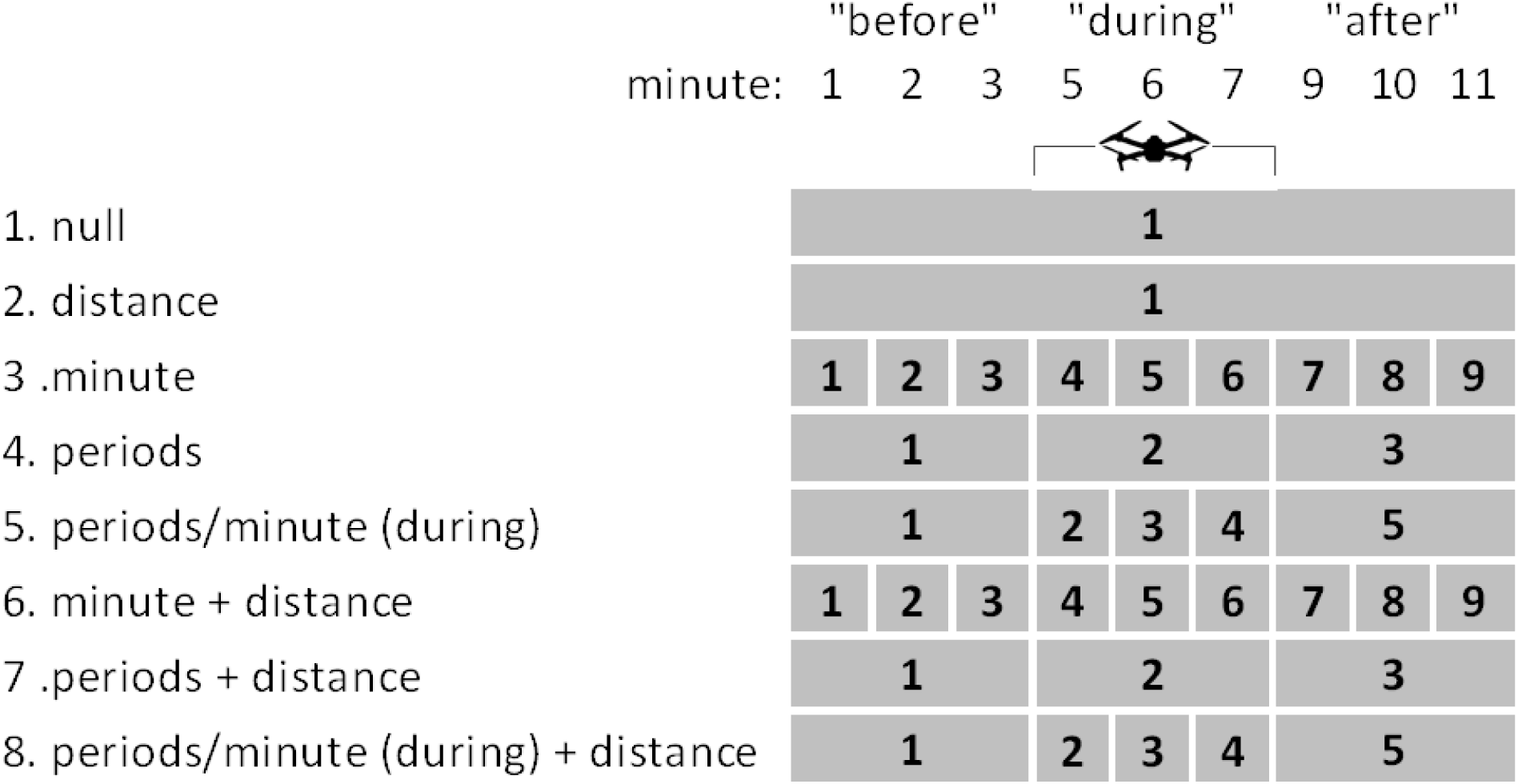
Gantt chart showing models used to test whether bird song output is related to the drone’s presence. All models include individual bird as a random effect. The null model and model 2 do not include time effects. Time effects of other models shown by numbered sequences. Note that minutes 4 (drone approaching) and 8 (drone departing) of the experiment were not included in the statistical analysis.

We used the same eight model structures illustrated in Figure 2 to test whether the singing rate (number of song bouts per minute) of each bird varied according to time period, this time with a Poisson error distribution. The same Poisson model was used to test whether the Northern Cardinal’s “tik” contact/alarm call showed time dependent patterns consistent with a response to the drone flight.

## RESULTS

### Probability of singing bird detection

We identified 8,303 song bouts across the seven study species, which we initially estimated to belong to 467 different individual birds (Table 1). However, these values are likely overestimates because ∼50% of the song bouts were localized to distances over 100 m from the array centers, and hence duplication of individuals between adjacent arrays was likely. To provide a more accurate analysis, we excluded song bouts that were localized to distances greater than 100m from the array centers. Further, many song bout detections were either not localizable (typically because they were not clear enough in all 4 spectrograms), or had large localization errors (greater than 5 ms) and were excluded from data analysis.

**Table 1.** Total number of song bouts detected, and number of bouts localized within 100 m and 50 m of the array centers, and number of individual birds within the 50 m radius.

|  | Total<br>song bout<br>detected | Localized song bouts |  | Birds within<br>50-m of array<br>centers |
| --- | --- | --- | --- | --- |
|  |  | Within 100-m<br>of array centers | Within 50-m of<br>array centers |  |
| American Robin | 1,597 | 858 | 331 | 18 |
| Yellow Warbler | 1,643 | 759 | 529 | 42 |
| Common Yellowthroat | 734 | 382 | 137 | 13 |
| Field Sparrow | 1,388 | 532 | 388 | 31 |
| Song Sparrow | 1,661 | 963 | 470 | 42 |
| Indigo Bunting | 611 | 357 | 241 | 14 |
| Northern Cardinal | 669 | 294 | 189 | 24 |
| all seven species | 8,303 | 4,145 | 2,285 | 184 |

The number of singing birds detected 0 m to 50 m from the array center was less variable over time than those detected 50 m to 100 m from center (Figure 3). Song bout detections showed a decrease in the three minutes when the drone was overhead, but most of this decline was of more distant birds, while the number of detections within 50 m was less variable time (Figure 3). However, there were clear differences among the seven study species. For three species: Common Yellowthroat, Song Sparrow, and Indigo Bunting, the patterns did not suggest a significant response to the drone. Northern Cardinal detections increased during the drone flight, and the increased song output continued 4-minutes after the drone had departed (Figure 3). There were overall reductions in the number of singing American Robins, Yellow Warblers, and Field Sparrows when the drone was overhead, but most of this apparent decrease was of birds more than 50 m from the array center (Figure 3), hence further away from both the drone and ARUs.

**Figure 3.**
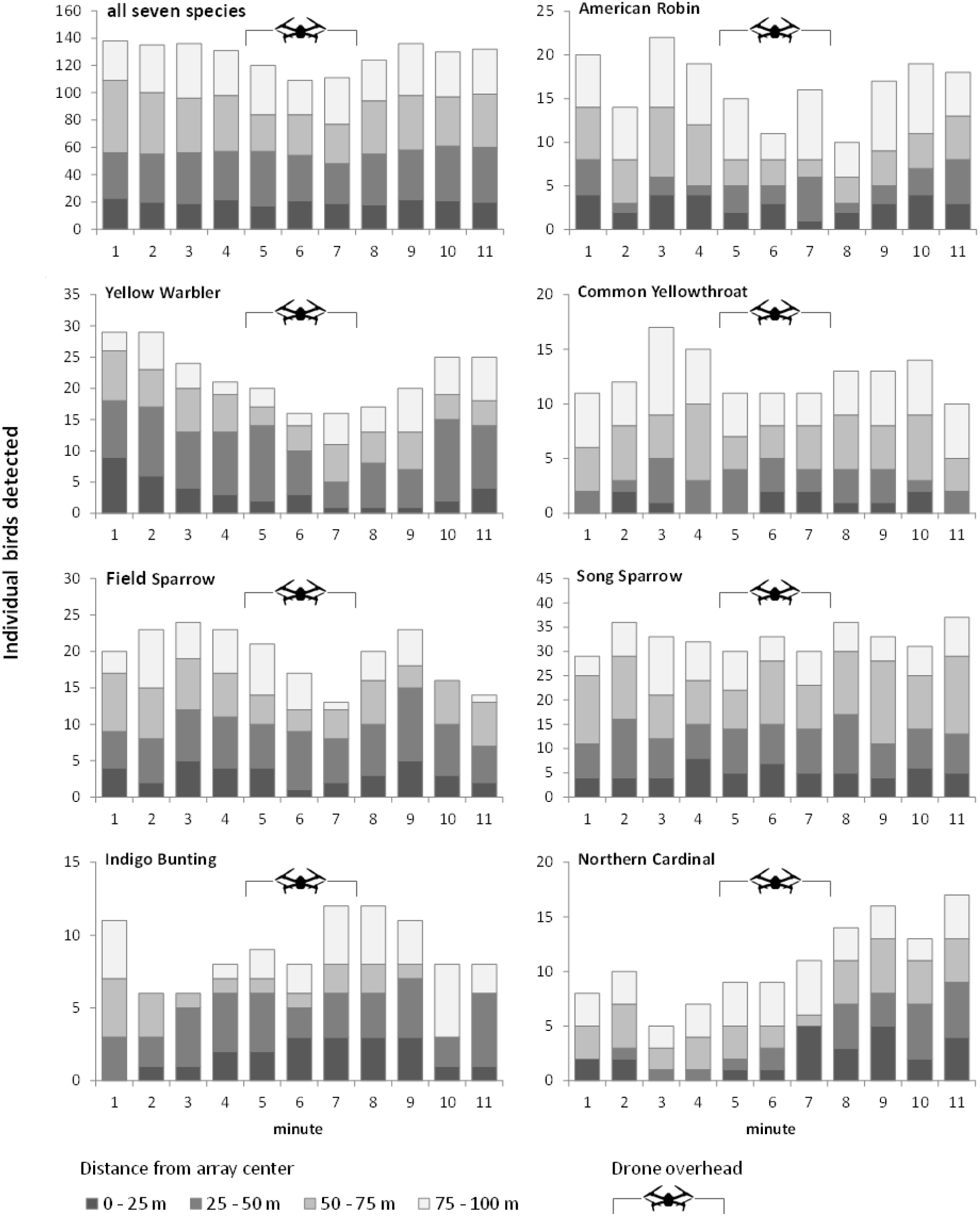
Number of individual signing birds detected by minutes and within four radial 25 meter distance bands from the array center.

Song bout detections showed a decrease in the three minutes when the drone was overhead, but most of this decline was of more distant birds, while the number of detections within 50 m was less variable over time (Figure 3). However, there were clear differences among the seven study species. For three species: Common Yellowthroat, Song Sparrow, and Indigo Burning, the patterns did not suggest a significant response to the drone. Detections of American Robin, Yellow Warbler, and Field Sparrow all showed a decline when the drone was overhead, while Northern Cardinal detections increased during the drone flight, and the increased song output persisted 4-minutes after the drone had departed (Figure 3).

Because most of the decline in song detections during the drone flight was of more distant birds (50-100 m from array centers; Figure 3), we assume that this was due to masking of songs by the drone noise on the recordings; it is unlikely that distant birds responded to the drone more than those closer to it. For this reason, we restricted our statistical analysis to the 2,285 song bouts of birds localized to within 50 m of the array centers (mean distance across bouts), which we attributed to 184 individual birds (Table 1). Yellow Warbler and Song Sparrow were the most abundant species, followed by Field Sparrow; all three species averaged more than one singing bird per array (within 50 m radius).

For five of the seven species, the null model (no time effects) GLMM had the best fit (lowest AIC), indicating no evidence for an effect of the drone on song detection within a 50 m radius (Table S1). The null model was also the best model when data for all seven species were combined. The two exceptions were the Yellow Warbler and the Northern Cardinal. For the Yellow Warbler, model 5 was the best fit, which indicates that song output was depressed when the drone was overhead and that the effect increased over time, such that there was a non-significant (Wald-test; p=0.311) 34% decrease in detections in the first minute, a non-significant 54% (P=0.035) decrease in the second minute, and a highly significant 81% (P=0.001) decrease in the third minute, when compared with the period before the drone flight (Table 2). Song detections remained 51% lower after the drone had departed (P=0.013). For the Northern Cardinal, the drone effect was also significant, but the results were the opposite of those for the Yellow Warbler—instead, cardinal song detections increased by a non-significant 46% (P=0.45) when the drone was overhead, and increased almost five-fold (P<0.001) after the drone had departed (Table 2).

**Table 2.** Results of GLMMs for models showing significant effects of the drone on detections of birds within 50 m of the array centers. Estimates (Est) are odd-ratios relative the first time period (either minute 1, or period 1). UL and UL are lower and upper confidence limits, and p-values are from Wald tests.

| Yellow Warbler |  |  |  |  |  |
| --- | --- | --- | --- | --- | --- |
|  |  | Est | LL | UL | P |
| Intercept |  | 0.665 | 0.407 | 1.087 | 0.103 |
| During | Minute 5 | 0.662 | 0.297 | 1.472 | 0.311 |
|  | Minute 6 | 0.438 | 0.189 | 1.017 | 0.054 |
|  | Minute 7 | 0.189 | 0.066 | 0.495 | 0.001 |
| After | Minutes 9-11 | 0.483 | 0.271 | 0.860 | 0.013 |

| Northern Cardinal |  |  |  |  |  |
| --- | --- | --- | --- | --- | --- |
|  |  | Est | LL | UL | P |
| Intercept |  | 0.111 | 0.050 | 0.248 | <0.001 |
| During | Minutes 5-6 | 1.462 | 0.544 | 3.925 | 0.450 |
| After | Minutes 9-11 | 4.888 | 1.977 | 12.084 | <0.001 |

### Singing rate

The null model (no time effects) was the best model for singing-rate for four species: Yellow Warbler, Field Sparrow, Song Sparrow, and Common Yellowthroat (Table S2). For the American Robin, model 7 was the best fit, but none of the estimates for time periods were significant (Table 3). For the Indigo Bunting, model 4 was the best fit, but the AIC was only marginally lower than that of the null model, and none of the estimates for time periods were significant (Table 3). We conclude that there was no evidence that singing-rates changed in response to the drone for six of the seven species. Once again, the results for Northern Cardinal were anomalous with a significant drop in singing-rate in minute 6 (P=0.019; Table 3), although once again, the time-effect model (5) did not appreciably improve model fit over the null model (Table S2).

**Table 3.** Results of GLMMs for models showing significant effects of the drone on singing rates of birds within 50 m of the array centers. Estimates (Est) are odd-ratios relative the first time period (either minute 1, or period 1). UL and UL are lower and upper confidence limits, and p-values are from Wald tests.

|  |  | Est | LL | UL | P |
| --- | --- | --- | --- | --- | --- |
| Intercept |  | 6.124 | 4.274 | 8.774 | <0.001 |
| During | Minutes 5-7 | 0.847 | 0.605 | 1.188 | 0.337 |
| After | Minutes 9-11 | 1.240 | 0.922 | 1.669 | 0.155 |
| distance |  | 0.984 | 0.972 | 0.995 | 0.005 |

Indigo Bunting
|  |  | Est | LL | Est | LL |
| --- | --- | --- | --- | --- | --- |
| Intercept |  | 4.846 | 3.786 | 6.204 | <0.001 |
| During | Minutes 5-7 | 0.711 | 0.501 | 1.009 | 0.056 |
| After | Minutes 9-11 | 0.728 | 0.511 | 1.037 | 0.079 |

Northern Cardinal
|  |  | Est | LL | Est | LL |
| --- | --- | --- | --- | --- | --- |
| Intercept |  | 3.792 | 2.494 | 5.764 | <0.001 |
| During | { Minute 5 | 0.759 | 0.351 | 1.640 | 0.483 |
|  | Minute 6 | 0.231 | 0.068 | 0.783 | 0.019 |
|  | Minute 7 | 0.942 | 0.506 | 1.754 | 0.851 |
| After | Minutes 9-11 | 0.782 | 0.491 | 1.247 | 0.302 |

all seven species
|  |  | Est | LL | Est | LL |
| --- | --- | --- | --- | --- | --- |
| Intercept |  | 3.736 | 3.200 | 4.363 | <0.001 |
| During | Minutes 5-7 | 0.907 | 0.804 | 1.024 | 0.115 |
| After | Minutes 9-11 | 0.968 | 0.862 | 1.087 | 0.586 |
| distance |  | 0.996 | 0.991 | 1.000 | 0.055 |

### Cardinal calls

A total of 1,148 Northern Cardinal “tik” calls were detected on recordings from ARU “A”, these detections coming from 23 of the 30 arrays. The calling rate increase from an average of 3.21/minute (se 1.39) during minutes 1 to 4 to 5.96/minute (se 1.11) in minutes 4 to 9, then declined to 2.60/minute (se 0.79) in the last two minutes of the experiment (Figure 4). Calling rates were significantly higher in minutes 5 thru 9 compared with minute 1 (Wald Tests, P<0.001). It should be noted that there appeared to be an inflection point in call rate at about 3 minute 30 second, which is around the time that the drone was first audible in the recording as it approached the array (Figure 4).

**Figure 4.**
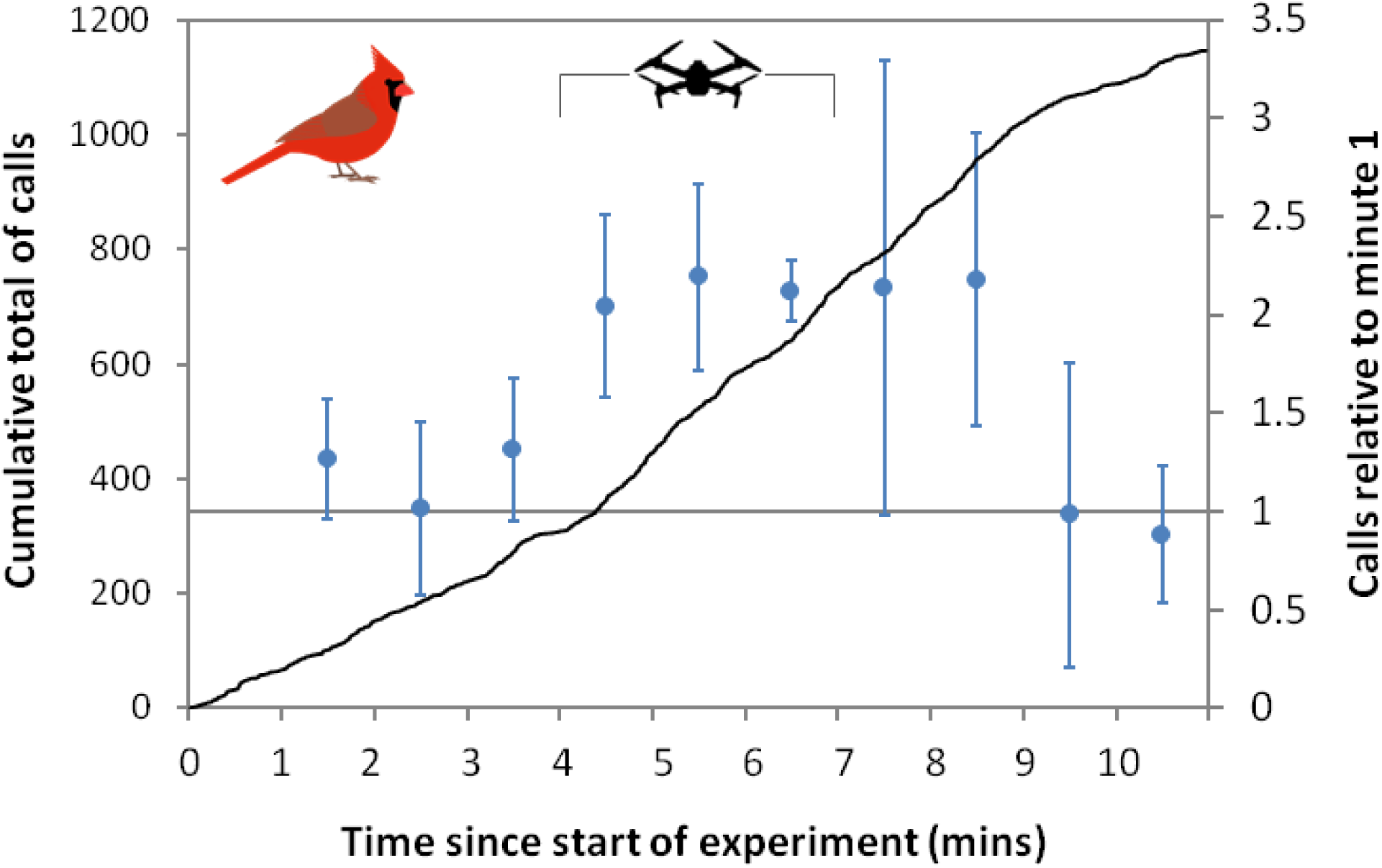
Calling rate of Northern Cardinals from a single ARU across all 30 arrays. Black line shows the cumulative total of calls and blue circles show GLMM coefficients for each 1-minute interval, relative to minute 1, with 95% confidence limits.

## DISCUSSION

Drawing on multiple lines of evidence, we conclude that overall, a drone hovering at 48 m a.g.l. for a 3-minute duration, causes little change in songbird output in the area below the drone, within a 50-m radius. However, there were significant impacts of the drone on the detection of songs of two species: the Yellow Warbler, which saw a substantial drop in song detection while the drone was overhead, and the Northern Cardinal, which saw a significant increase in song detections, especially after the drone’s departure. The pronounced increase in Northern Cardinal contact/alarm calls during and following the drone flights provides further evidence that the vocal behavior of that species was affected by the drone’s presence.

The fact that we found that responses to the drone were varied among a small group of passerines is intriguing. Differential responses to anthropogenic noise disturbance have been detected by others (Francis, Ortega and Cruz, 2009; McClure *et al*., 2013; Roca *et al*., 2016). The differences have, in part, been attributed to whether the vocalization frequency of a bird species overlaps with the frequency of the noise disturbance (Francis, Ortega and Cruz, 2009), presumably because birds would not waste energy vocalizing if it was likely to be masked by other noise. However, of our seven study species, two have relative low frequency songs (American Robin and Northern Cardinal; peak sound pressure 2 to 4 kHz), Field Sparrows are slightly higher (peak sound pressure 3 to 5 kHz) and the others have higher frequency songs (peak sound pressure 4 to 8 kHz) (Pieplow, 2017), so we find no discernible association between sound frequency and effects of the drone. Being a small vehicle, the noise emitted by a DJI Mavic Pro is of higher frequencies than larger drones, and with a broad spectrum (Miljkovic, 2018), which overlaps with the songs of all species included in our study. The increased singing and calling of cardinals is indicative of their agitation at the drone noise. The Northern Cardinal is a species that has been show to increase its song frequency in response to anthropogenic noise (Dowling, Luther and Marra, 2012; Seger-Fullam, Rodewald and Soha, 2012).

It should be noted that we chose the DJI Mavic Pro for this study because of its suitability for conducting aerial bioacoustics surveys: it is small and therefore quieter than the larger vehicles that are typically used in wildlife research. Although we don’t have our own measurement of the drone noise from the ground, the manufacturer reports sound pressure (SPLa) of 70 dB at 1 meter (per Valle and Scarton 2019), which would attenuate to around 36 dB at ground-level directly below a drone hovering at 48 m, and around 33 dB at ground-level at the edge of the 50 m radius. In a review of anthropogenic disturbance affects, it was found that impacts on bird song, abundance, reproduction, and stress hormone levels were typically found when anthropogenic noise levels exceeded 45 dB SPL (Shannon *et al*., 2016). Therefore, while not wishing to downplay the noise pollution emitted from small drones, the noise levels that birds were subjected to in our experiment were modest in comparison to some other sources of anthropogenic noise. However, the relatively high frequency of drone noise compared to most forms of anthropogenic noise pollution, which is loudest below 2 kHz (Kight, Saha and Swaddle, 2012), results in more overlap with the frequency of most bird song, and hence it could be that drone noise disturbance to birds is a function of its frequency, rather than loudness.

It has previously been shown that prolonged exposure to drones has an increased disturbance effect (Mulero-Pázmány *et al*., 2017). Our results support this with impacts most evident after 2 minutes of exposure for both the Yellow Warbler and Northern Cardinal. If drones are to be used for bioacoustics surveys for birds, we recommend that very short duration counts are used. Not only does this reduce disturbance, but it also allows for multiple counts to be made on a single drone battery. If short duration counts are considered inadequate, for example if the probability of detecting a bird is too low, an alternative approach would be to do repeated counts over the duration of a day, or perhaps over several days. This would then provide a sampling framework that was amenable to estimating species presence or abundance using occupancy models (Royle and Nichols, 2003; Hayes and Monfils, 2015) or spatially replicated N-mixture models (Royle, 2004). An added advantage of using drones for aerial bioacoustics surveys under those circumstances is that they are highly replicable, both in terms of location and duration (due the ability to precisely program missions), and because audio recordings provide a permanent record of bird song that can be analyzed by multiple observers after the fact (Shonfield and Bayne, 2017).

A limiting factor in our study is that despite the very large number of song bouts analyzed, sample sizes for the number of individual birds for some of our study species were small, which may have reduced the power to detect more modest responses. However, given the fact that many of the birds tracked were found to sing throughout the time that drone was overhead, and that there was no evidence that singing-rates declined for six of our seven species, we can be certain that effects were very limited for five of the seven species. It should also be noted that most of the species in our study sing from relatively low vegetation (at least at our study site, pers. obs.); the Indigo Bunting being an exception. Therefore, we can’t generalize our results to species that sing from more elevated perches, which would be closer to the drone and perhaps more susceptible to disturbance. Clearly, further studies of the impacts of drone on bird singing behavior are merited to see whether our results hold for a wider range of species, and different habitat types.

Measurement error, especially with respect to the sound localization process could also have impacted our results. Limiting the statistical analysis to within 50 m of the array center will have reduced localization errors because errors increase with distance from the array (Campbell and Francis, 2013; Wilson *et al*., 2014), but even so, we should expect that errors of 5 to 10 m were commonplace, even if the mean error was small (Wilson *et al*., 2014). Given that our study did not aim to estimate populations densities—a common application of sound localization—the effects of these errors on our findings may be limited to whether or not individual birds were included in the statistical analysis (i.e. whether were they <50 m from array center). We do not think that erroneous inclusion or exclusion of a few individuals would be sufficient to change our findings, especially as those marginal birds are furthest from the drone, and hence the least likely to be affected by its presence.

It is possible that our ability to individually identify birds based on their song pattern and location was not always accurate. Anecdotally, we observed that Yellow Warblers may have shown song-pattern plasticity (Lowther *et al*., 1999), which could potentially have resulted in us over-estimating the number of different birds. However, because we were using sound localization to track birds in space and time, we think cases of confusion due to individual bird song-switching would be limited to a few individuals at the most.

Our study focused on assessing the effects of drone noise on bird song for a specific vehicles type and altitude, which was due to our interest in using drones for aerial bioacoustics. However, our results have more general interest for those concerned about disturbance effects of drones on wildlife, and to our knowledge, ours is the first study to focus specifically on how drones affect bird vocalizing behavior. A survey of UAS survey altitudes, focusing on wildlife applications, shows that 50 m is frequently used for small UASs (Sherman and Chatowsky in. prep.).

That fact that we did find some bird behavioral responses to the drone in our study gives us reason for pause with respect to using drones for airborne bioacoustics. However, there has been a rapid improvement in technology to reduce unwanted drone noise, even in the time since our study. More recent DJI Mavic models are substantially quieter, with refined propeller shapes reportedly reducing sound pressure by 4 dB, which represent about 60% reduction in noise (Valle and Scarton, 2019). With the advent of quieter drones, we hope that concerns about noise pollution impacts on wildlife will diminish, but we urge for continued vigilance with regards to the behavioral impacts of using drones to study wildlife.

## Ethics Statement

Our experiments were approved by the Gettysburg College Institutional Animal Care and Use Committee (IACUC). The drone was operated by the US Federal Aviation Authority (FAA) Part 107 license holder A.M. Wilson, and in accordance with FAA guidelines.

## Acknowledgements

We thank the Pennsylvania Game Commission for permitting us to use State Game Lands 249 for the experiments. State Game Lands 249 is on the lands of the Susquehannock people. Gettysburg College students Taylor Derick, Arthur Garst, and Samantha Trueman helped with data analysis. The research was funding by the Gettysburg College Cross-Disciplinary Science Institute (X-SIG).

## Supplementary Materials

**Table S1.**
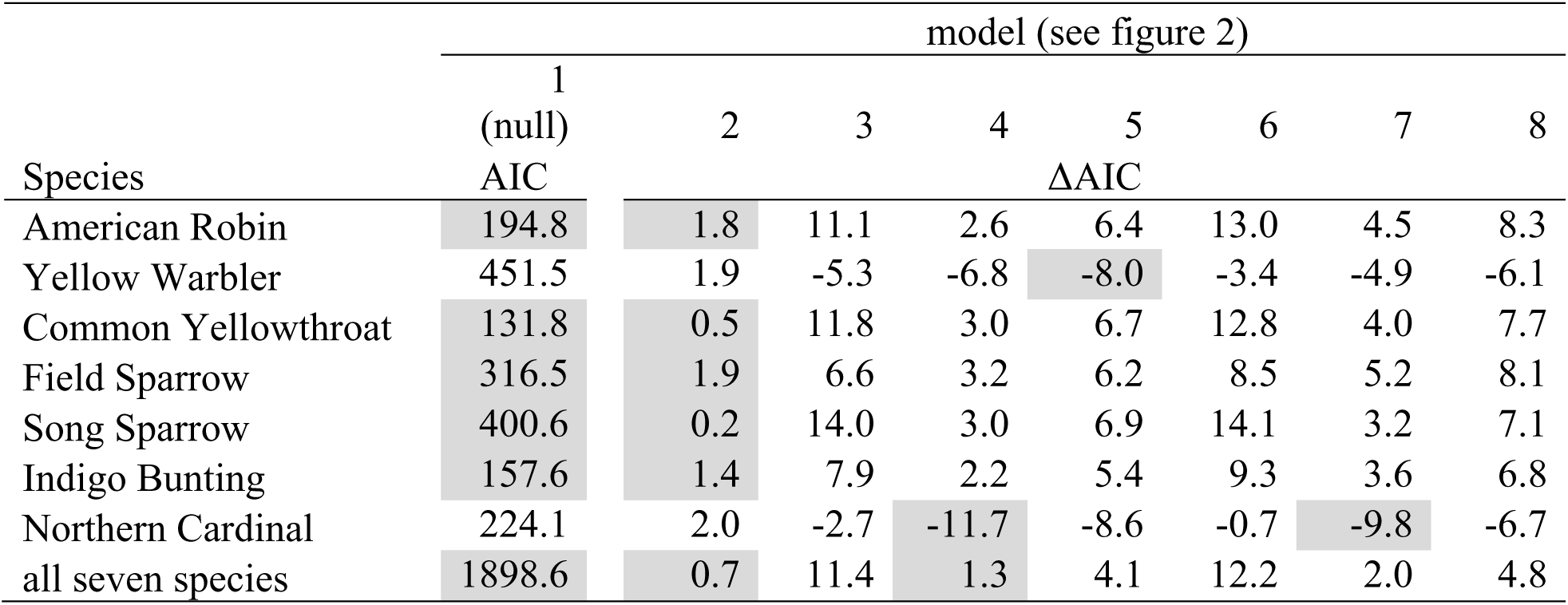
AIC values for the eight models for each species, for probability of detection. ΔAIC is compared with the null model (model 1). Gray shading indicates the lowest AIC (or joint lowest if ΔAIC less than 2).

| Species | model (see figure 2) |  |  |  |  |  |  |  |
| --- | --- | --- | --- | --- | --- | --- | --- | --- |
|  | 1 | 2 | 3 | 4 | 5 | 6 | 7 | 8 |
| | (null)<br>AIC | | | | ΔAIC | | | |
| American Robin | 194.8 | 1.8 | 11.1 | 2.6 | 6.4 | 13.0 | 4.5 | 8.3 |
| Yellow Warbler | 451.5 | 1.9 | -5.3 | -6.8 | -8.0 | -3.4 | -4.9 | -6.1 |
| Common Yellowthroat | 131.8 | 0.5 | 11.8 | 3.0 | 6.7 | 12.8 | 4.0 | 7.7 |
| Field Sparrow | 316.5 | 1.9 | 6.6 | 3.2 | 6.2 | 8.5 | 5.2 | 8.1 |
| Song Sparrow | 400.6 | 0.2 | 14.0 | 3.0 | 6.9 | 14.1 | 3.2 | 7.1 |
| Indigo Bunting | 157.6 | 1.4 | 7.9 | 2.2 | 5.4 | 9.3 | 3.6 | 6.8 |
| Northern Cardinal | 224.1 | 2.0 | -2.7 | -11.7 | -8.6 | -0.7 | -9.8 | -6.7 |
| all seven species | 1898.6 | 0.7 | 11.4 | 1.3 | 4.1 | 12.2 | 2.0 | 4.8 |

**Table S2.** AIC values for the eight models for each species, for singing-rate. ΔAIC is compared with the null model (model 1). Gray shading indicates the lowest AIC (or joint lowest if ΔAIC less than 2).

| Species | model (see figure 2) |  |  |  |  |  |  |  |
| --- | --- | --- | --- | --- | --- | --- | --- | --- |
|  | 1 | 2 | 3 | 4 | 5 | 6 | 7 | 8 |
| | (null)<br>AIC | ΔAIC | | | | | | |
| American Robin | 264.1 | -4.0 | 7.2 | -0.4 | 1.8 | 2.5 | -5.5 | -4.0 |
| Yellow Warbler | 492.2 | 2.5 | 10.8 | 2.7 | 6.5 | 12.7 | 4.6 | 8.5 |
| Common Yellowthroat | 146.4 | 1.6 | 3.6 | 3.8 | 6.0 | 5.6 | 5.3 | 7.7 |
| Field Sparrow | 383.3 | 1.6 | 15.1 | 3.6 | 7.3 | 16.7 | 5.2 | 9.0 |
| Song Sparrow | 468.1 | 2.0 | 13.7 | 3.3 | 6.7 | 15.7 | 5.2 | 8.6 |
| Indigo Bunting | 202.5 | 1.7 | 10.3 | -0.1 | 3.4 | 12.2 | 1.8 | 5.4 |
| Northern Cardinal | 193.0 | 1.7 | 6.6 | 2.3 | -0.3 | 8.5 | 4.1 | 1.6 |
| all seven species | 2152.2 | -1.8 | 7.7 | 1.1 | 4.0 | 6.0 | -0.4 | 2.5 |

## LITERATURE CITED

Afán, I., Máñez, M. and Díaz-Delgado, R. (2018) ‘Drone Monitoring of Breeding Waterbird Populations: The Case of the Glossy Ibis’, Drones, 2(4), p. 42. doi: 10.3390/drones2040042.

Barnas, A. F. et al. (2019) ‘A comparison of drone imagery and ground-based methods for estimating the extent of habitat destruction by lesser snow geese (Anser caerulescens caerulescens) in La Pérouse Bay’, PloS one. Public Library of Science, 14(8), pp. e0217049–e0217049. doi: 10.1371/journal.pone.0217049.

Bates, D. et al. (2015) ‘Fitting linear mixed-effects models using lme4’, Journal of Statistical Software, 67(1), pp. 0–48. doi: 10.18637/jss.v067.i01.

Borrelle, S. B. and Fletcher, A. T. (2017) ‘Will drones reduce investigator disturbance to surface-nesting birds?’, Marine Ornithology, 45(1), pp. 89–94.

Campbell, M. and Francis, C. M. (2013) ‘Using Stereo-Microphones to Evaluate Observer Variation in North American Breeding Bird Survey Point Counts’, http://dx.doi.org/10.1525/auk.2011.10005. University of California Press.

Christiansen, F. et al. (2016) ‘Noninvasive unmanned aerial vehicle provides estimates of the energetic cost of reproduction in humpback whales’, Ecosphere. John Wiley & Sons, Ltd, 7(10), p. e01468. doi: 10.1002/ecs2.1468.

Dowling, J. L., Luther, D. A. and Marra, P. P. (2012) ‘Comparative effects of urban development and anthropogenic noise on bird songs’, Behavioral Ecology, 23(1), pp. 201–209. doi: 10.1093/beheco/arr176.

Francis, C. D., Ortega, C. P. and Cruz, A. (2009) ‘Noise Pollution Changes Avian Communities and Species Interactions’, Current Biology, 19(16), pp. 1415–1419. doi: 10.1016/j.cub.2009.06.052.

Hayes, D. B. and Monfils, M. J. (2015) ‘Occupancy modeling of bird point counts: Implications of mobile animals’, Journal of Wildlife Management, 79(8), pp. 1361–1368. doi: 10.1002/jwmg.943.

Inger, R. et al. (2015) ‘Common European birds are declining rapidly while less abundant species’ numbers are rising’, Ecology Letters. John Wiley & Sons, Ltd, 18(1), pp. 28–36. doi: 10.1111/ele.12387.

Kelaher, B. P. et al. (2020) ‘Assessing variation in assemblages of large marine fauna off ocean beaches using drones’, Marine and Freshwater Research, 71(1), pp. 68–77.

Kight, C. R., Saha, M. S. and Swaddle, J. P. (2012) ‘Anthropogenic noise is associated with reductions in the productivity of breeding Eastern Bluebirds (Sialia sialis)’, Ecological Applications, 22(7), pp. 1989–1996. doi: 10.1890/12-0133.1.

Kleist, N. J. et al. (2016) ‘Sound settlement: Noise surpasses land cover in explaining breeding habitat selection of secondary cavity nesting birds’, Ecological Applications. doi: 10.1002/eap.1437.

Kloepper, L. N. and Kinniry, M. (2018) ‘Recording animal vocalizations from a UAV: Bat echolocation during roost re-entry’, Scientific Reports. Springer US, 8(1), pp. 8–13. doi: 10.1038/s41598-018-26122-z.

Lowther, P. E. et al. (1999) ‘Yellow Warbler (Dendroica petechia)’, The Birds of North America Online. edited by A. Poole and F. Gill. doi: 10.2173/bna.454.

McClure, C. J. W. et al. (2013) ‘An experimental investigation into the effects of traffic noise on distributions of birds: avoiding the phantom road.’, Proceedings. Biological sciences / The Royal Society. The Royal Society, 280(1773),. 20132290. doi: 10.1098/rspb.2013.2290.

Miljkovic, D. (2018) ‘Methods for attenuation of unmanned aerial vehicle noise’, 2018 41st International Convention on Information and Communication Technology, Electronics and Microelectronics, MIPRO 2018 - Proceedings, (May), pp. 914–919. doi: 10.23919/MIPRO.2018.8400169.

Mulero-Pázmány, M. et al. (2017) ‘Unmanned aircraft systems as a new source of disturbance for wildlife: A systematic review’, PLOS ONE. edited by A. Margalida. Public Library of Science, 12(6), p. e0178448. doi: 10.1371/journal.pone.0178448.

Musgrove, A. et al. (2013) ‘Population estimates of birds in Great Britain and the UK’, British Birds, 106(4), pp. 231–232.

Pieplow, N. (2017) Peterson Field Guide to Bird Sounds of Eastern North Americae. New York, New York, USA: Houghton Mifflin Harcour.

Ratcliffe, N. et al. (2015) ‘A protocol for the aerial survey of penguin colonies using UAVs1’, http://dx.doi.org/10.1139/juvs-2015-0006. NRC Research Press http://www.nrcresearchpress.com.

Roca, I. T. et al. (2016) ‘Shifting song frequencies in response to anthropogenic noise: a meta-analysis on birds and anurans’, Behavioral Ecology, 27(5), pp. 1269–1274. doi: 10.1093/beheco/arw060.

Rosenberg, K. V et al. (2019) ‘Decline of the North American avifauna’, Science, 366(6461), pp. 120 LP – 124. doi: 10.1126/science.aaw1313.

Royle, J. A. (2004) ‘N-mixture models for estimating population size from spatially replicated counts’, Biometrics, 60(1), pp. 108–115. Available at: http://pubs.er.usgs.gov/publication/5224798.

Royle, J. A. and Nichols, J. D. (2003) ‘Estimating abundance from repeated presence-absence data or point counts’, Ecology, 84(3), pp. 777–790. doi: 10.1890/0012-9658(2003)084[0777:EAFRPA]2.0.CO;2.

Scholten, C. N. et al. (2019) ‘Real-time thermal imagery from an unmanned aerial vehicle can locate ground nests of a grassland songbird at rates similar to traditional methods’, Biological Conservation, 233, pp. 241–246. doi: https://doi.org/10.1016/j.biocon.2019.03.001.

Seger-Fullam, K. D., Rodewald, A. D. and Soha, Ji. A. (2012) ‘Urban noise predicts song frequency in Northern Cardinals and American Robins’, Bioacoustics. Taylor & Francis Group, 20(3), pp. 267–276. doi: 10.1080/09524622.2011.9753650.

Shannon, G. et al. (2016) ‘A synthesis of two decades of research documenting the effects of noise on wildlife’, Biological Reviews, 91(4), pp. 982–1005. doi: 10.1111/brv.12207.

Shonfield, J. and Bayne, E. M. (2017) ‘Autonomous recording units in avian ecological research: current use and future applications’, Avian Conservation and Ecology, 12(1). doi: 10.5751/ace-00974-120114.

Valle, R. G. and Scarton, F. (2019) ‘Effectiveness, efficiency, and safety of censusing eurasian oystercatchers haematopus ostralegus by unmanned aircraft’, Marine Ornithology, 47(1), pp. 81–87.

Vermeulen, C. et al. (2013) ‘Unmanned aerial survey of elephants’, PloS one. 2013/02/06. Public Library of Science, 8(2), pp. e54700–e54700. doi: 10.1371/journal.pone.0054700.

Weissensteiner, M. H., Poelstra, J. W. and Wolf, J. B. W. (2015) ‘Low-budget ready-to-fly unmanned aerial vehicles: an effective tool for evaluating the nesting status of canopy-breeding bird species’, Journal of Avian Biology. Blackwell Publishing Ltd, 46(4), pp. 425–430. doi: 10.1111/jav.00619.

Wildlife Acoustics (2019) ‘Kaleidoscope Pro Analysis Software’. Boston, MA. Available at: https://www.wildlifeacoustics.com/products/kaleidoscope-pro/overview.

Wilson, A. M., Barr, J. and Zagorski, M. (2017) ‘The feasibility of counting songbirds using unmanned aerial vehicles’, The Auk. The American Ornithologists’ Union, 134(2), pp. 350–362. doi: 10.1642/auk-16-216.1.

Wilson, D. R. et al. (2014) ‘Sound Finder: a new software approach for localizing animals recorded with a microphone array’, Bioacoustics. Taylor & Francis, 23(2), pp. 99–112. doi: 10.1080/09524622.2013.827588.

Witczuk, J. et al. (2017) ‘Exploring the feasibility of unmanned aerial vehicles and thermal imaging for ungulate surveys in forests - preliminary results’, International Journal of Remote Sensing. Taylor & Francis, pp. 1–18. doi: 10.1080/01431161.2017.1390621.

